# Rapid genome functional annotation pipeline anchored to the House sparrow (*Passer domesticus*, Linnaeus 1758) genome reannotation

**DOI:** 10.1101/2023.01.27.525869

**Authors:** Melisa Eliana Magallanes-Alba, Agustín Barricalla, Natalia Rego, Antonio Brun, William H. Karasov, Enrique Caviedes-Vidal

## Abstract

House sparrow (*Passer domesticus*) is an important avian model for both laboratory and field-based studies of evolutionary genetics, development, neurobiology, physiology, behavior, and ecology. The current annotation of the *P. domesticus* genome is available at Ensembl Rapid Release site, which currently only focuses on gene set building. Here, we provide the first functional reannotation of the *P. domesticus* genome based on enrichment with intestinal Illumina RNA-Seq libraries. This revised annotation describes 38592 transcripts, compared to 23574 currently for Ensembl, and 14717 predicted protein-coding genes, with 96.4% complete Passeriformes lineage BUSCOs. A key improvement in this revised annotation is the definition of untranslated region (UTR) sequences, with 82.7% and 93.8% of transcripts containing 5’ and 3’ UTRs, respectively. Our reannotation highlights the benefits to genome annotation improvement when additional specific RNA-Seq data is available for analysis and rapid data throughput (>200 Mb h^−1^) is used.

## Introduction

Studies in the past decade herald the potential power of “omics” analyses to advance understanding of mechanistic bases underlying adaptations and differences in phenotypes within and across avian species (*1–4*). Good quality genome sequences and functional genome annotations represent essential resources for these studies. RNA sequencing (RNA-Seq) has been previously used to improve them, including: (i) correcting predicted gene structures (*5*); (ii) detecting new alternative splicing isoforms (*6*); and (iii) discovering new genes and new transcripts (*7, 8*). Features of RNA-Seq data available for genome annotation vary from case to case according to ontogenetic state, organ and tissue specificity, pathogens, and additional factors. Thus, opportunity for gene annotation improvement may always exist for those genes and transcripts expressed less widely, mainly in studies of specific tissues, ontogenetic states, or different experimental conditions.

We are using the omnivorous avian model *Passer domesticus* (house sparrows) to study physiological pathways and underlying mechanisms associated with nutritional flexibility (*9, 10*). The house sparrow is the most widely distributed wild bird in the world and somewhat unique among wild avian species in its close association with humans, not only in the agricultural environment, where presumably this association first evolved, but also in urban areas. Our functional studies are on intestine, and accurate gene identification is important considering the multiplicity of significant unique functions that rely on the single layer of epithelial cells, i.e., the enterocytes of the intestine of vertebrates in general (*11, 12*). To date, the genome of the house sparrow (GCA_001700915.1) was sequenced by Illumina HiSeq technology, and its current gene set annotation has been determined using Ensembl Rapid release (https://rapid.ensembl.org/Passer_domesticus_GCA_001700915.1/Info/Index), which is based on the Ensembl Genebuild Method (*13*). The protein-coding annotation method includes an RNA-Seq pipeline as one of the main data sources for model generation, which in the case of the current annotation involved only one individual’s sample of Illumina RNA-seq data from only one tissue, muscle, to the target genome. Therefore, it is likely that a considerable proportion of genes are mis-annotated in the reference gene set. Indeed, this problem became apparent during our analysis of RNA-Seq samples from house sparrows of different ages feeding on different diets. The current Ensembl annotation revealed only 45.3% and 48.6% of its transcripts containing 5’ and 3’ untranslated regions (UTRs), respectively. Many studies highlight the importance of short open reading frames (ORFs) in 5’ UTRs in fine-tunning gene functions (*14*). 3’ UTR sequences are frequently needed to design experiments (*15*). Therefore, our work was aimed at generating an improved new genome annotation version with improved relevance to intestinal tissue. We used intestinal tissue expression data obtained from a previous experiment with wild-hatched house sparrow nestlings that were removed from nests at age 3 days old (d.o) and then raised in environmental chambers using a semisynthetic starch based diet (*16*). To achieve our goal, we first used Illumina technology for RNA-Seq to assess RNA expression from experimental samples and then the GeMoMa pipeline (*17*) to build the structural and functional annotation. An additional goal was to optimize the time to complete the de novo annotation using modest or low computing resources (ie., 20 CPUs and 100GB RAM). The GeMoMa-based pipeline proposed in this work used on our intestinal RNA-seq dataset proved to offer an adequate data resource to re-annotate the genome.

## Material and Methods

### Study site and sample collection

All samples were obtained from a previous experiment with house sparrow nestlings aimed at studying the phenotypic flexibility of the intestinal enzymes, maltase, sucrase and aminopeptidase-N (*16*). Briefly, three days post-hatch nestlings were collected from their wild nests and housed individually in environmental controlled chambers in our laboratory. At arrival birds were fed a high starch-low casein content diet and all 5 individual birds fed for 6 days continuously on this diet were used in this work (Rott et al. 2017). Nestlings were euthanized with CO_2_ and dissected to remove the small intestine, and medial regions were preserved in RNAlater (Invitrogen; Carlsbad, California, USA).

### RNA extraction, quantification, and integrity control

Total RNA was isolated from frozen tissue using the PureLink™ RNA Mini Kit (Invitrogen; Carlsbad, California, USA), according to the manufacturer’s instructions. Before all the procedures we decontaminated equipment, benchtops, glassware and plasticware using RNase AWAY™ Surface Decontaminant (Thermo Scientific; Waltham, MA, USA). Total RNAs were submitted to the University of Wisconsin-Madison Biotechnology Center for quantification. All samples were first assayed for purity and integrity using a NanoDropOne Spectrophotometer and an Agilent 2100 BioAnalyzer.

### Construction and Sequencing of Directional Libraries

The Illumina^®^ TruSeq^®^ Stranded mRNA Sample Preparation kit (Illumina Inc., San Diego, California, USA) was used to construct the libraries. For each library preparation, mRNA was purified from 1000 ng total RNA using poly-T oligo-attached magnetic beads. Subsequently, each poly-A enriched sample was fragmented using divalent cations under elevated temperature. Fragmented RNA was synthesized into double-stranded cDNA using SuperScript II Reverse Transcriptase (Invitrogen, Carlsbad, California, USA) and random primers for first strand cDNA synthesis followed by second strand synthesis using DNA Polymerase I and RNAse H for removal of mRNA. Double-stranded cDNA was purified by paramagnetic beads (Agencourt AMPure XP beads, Beckman Coulter). The cDNA products were incubated with Klenow DNA Polymerase to add an ‘A’ base (Adenine) to the 3’ end of the blunt DNA fragments. DNA fragments were ligated to Illumina unique dual adapters, which have a single ‘T’ base (Thymine) overhang at their 3’end. The adapter-ligated DNA products were purified by paramagnetic beads. Adapter ligated DNA was amplified in a Linker Mediated PCR reaction (LM-PCR) for 10 cycles using PhusionTM DNA Polymerase and Illumina’s PE genomic DNA primer set and then purified by paramagnetic beads. Quality and quantity of the finished libraries were assessed on the ATTI Fragment Analyzer (Agilent Technologies, Inc., Santa Clara, CA, USA) and Qubit^®^ dsDNA HS Assay Kit (Invitrogen, Carlsbad, California, USA), respectively. Libraries were standardized to 2nM. Paired-end 2×150 bp sequencing was performed on an Illumina NovaSeq6000 sequencer. Raw reads were submitted to the National Center of Biotechnology Information (NCBI) to the Sequence Read Archive under the Bioproject: PRJNA785148.

### *Passer domesticus* genome annotation

The publicly available genome assembly for *P. domesticus* (GCA_001700915.1) was “cleaned” to remove some repetitive contigs that are contained in other scaffolds, sorted by length, and masked using Funannotate v1.8.1 (*18*).

The *P. domesticus* genome assembly was annotated using GeMoMa v.1.9 (*17*). The GeMoMaPipeline function was run to complete the full pipeline with a maximum intron size of 200 kb using as reference three close related species genomes, *Passer montanus* (GCA_014805655.1), *Pyrgilauda ruficollis* (GCA_017590135.1) and *Onychostruthus taczanowskii* (GCA_017590135.1). These genomes were downloaded from NCBI (*19*) and represent the closest phylogenetically related species available at present that share same genus or family, and their annotated transcriptomes exhibit high Passeriformes lineage completeness levels in BUSCO (99.2% in all cases). The intestinal RNA-seq data generated (Table S1 in Supplemental file 1), and also muscle RNA-seq data available on Ensembl (*20; https://ftp.ensembl.org/pub/rapid-release/species/Passer_domesticus/GCA_001700915.1/rnaseq/*), were aligned to the *P. domesticus* genome using HISAT2 v.2.2.1 (*21*), and the aligned bam file incorporated to the pipeline. To avoid any possible complications at the alignment step, the RNA-seq data were previously analyzed by Rcorrector (*22*) and TrimGalore (*23*) to correct sequencing errors and trim adapters, respectively. Since GeMoMa simply uses the mapped RNA-seq data to predict UTRs and identify introns and splice sites only in previously built gene models, missing any expressed genes absent in the chosen reference organisms. Therefore, the aligned short reads were assembled into transcripts using Stringtie v2.2.1 (*24*), and the GFF file generated was incorporated to the GeMoMaPipeline through its external annotation option.

The structural annotation obtained from GeMoMa was analyzed using BUSCO (*25*) to identify the conserved gene protein dataset of the Passeriformes lineage. Additionally, miniprot (*26*) was run to align protein to genome using as query the Passeriformes dataset, obtaining as a result a second structural annotation. Then, the structural annotations generated by both GeMoMa and miniprot were merged using the ‘agat_sp_merge_annotations.pl script’ (*27*).

To generate the functional annotations of the final set of protein-coding genes, each of the predicted protein sequences was searched against the InterPro protein database via InterProScan v5.57–9.0 (*28*). Then, functional annotations against the Carbohydrate-Active Enzymes (CAZy), Clusters of Orthologous Groups of proteins (COG), Gene Ontology (GO), and Kyoto Encyclopedia of Genes and Genomes (KEGG) databases were derived using EggNOG-mapper v2.1.9 (*29*). The results were incorporated to the annotation using the AGAT’s script: ‘agat_sp_manage_functional_annotation’ (*27*) (Figure 1).

**Figure 1.**
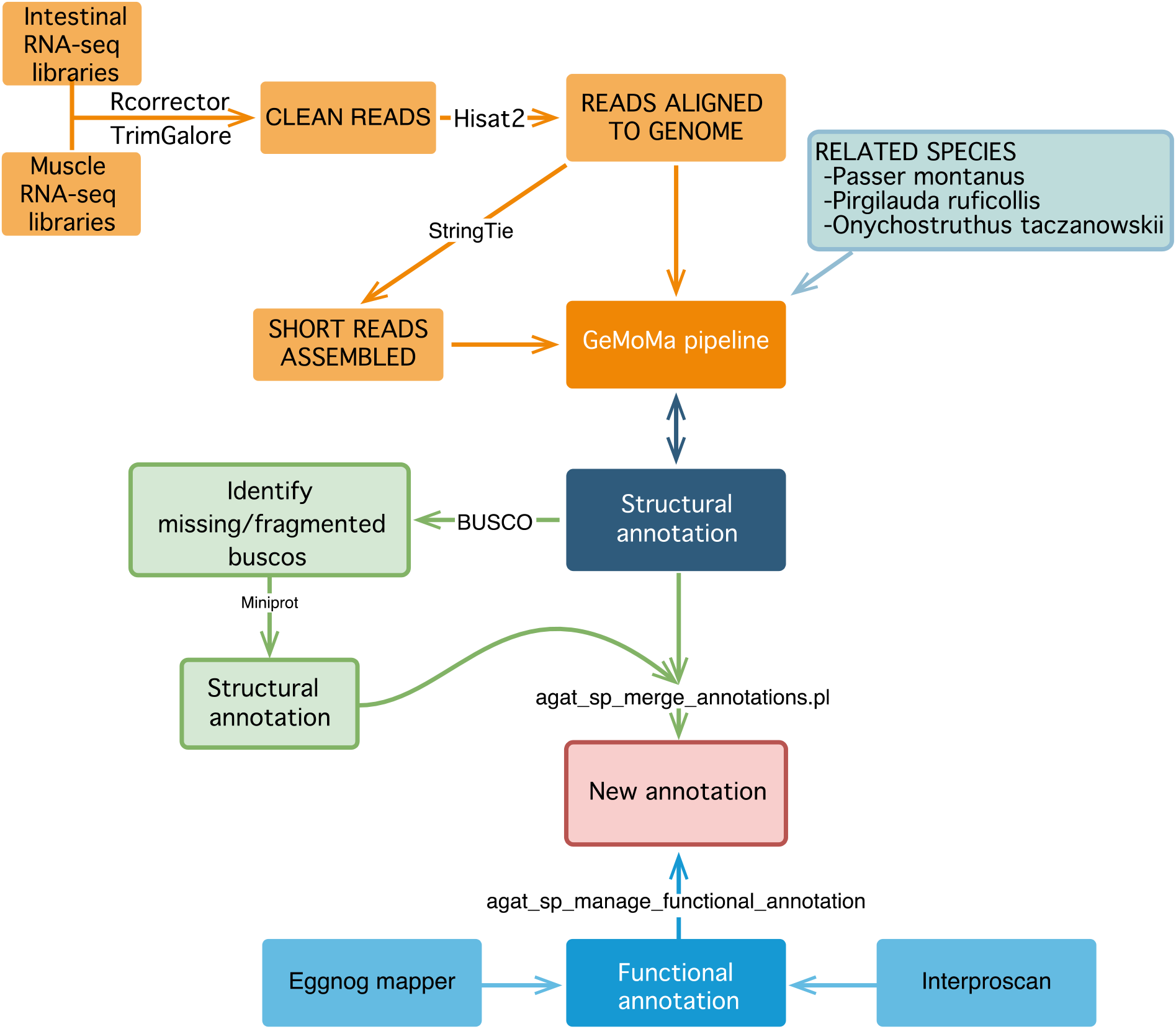
Annotation workflow for *P. domesticus* protein-coding genes.

### Comparison of annotations

The annotation summaries were generated using the ‘agat_sp_functional_statistics.pl script’ (*27*). To avoid a high proportion of duplicates, the longest isoform of each protein-coding gene was selected using the ‘agat_sp_keep_longest_isoform.pl’ (*27*), and BUSCO v5.3.2 (*30*) was used to assess the completeness of the genome and annotations based on the curated set of Passeriformes and Aves lineage specific single copy orthologs (from OrthoDB passeriformes_odb10 and aves_odb10 databases, *31*). Finally, Integrative Genomics Viewer (IGV) (*32*) was used for manual inspection of gene annotations across the genome.

Computing resources were provided by the University of Wisconsin - Madison Center for High Throughput Computing (CHTC) of the Department of Computer Sciences (University of Wisconsin-Madison, 53706 WI).

## Results and Discussion

### Reannotation of the *P. domesticus* genome

In this study, we created an updated gene annotation of the *P. domesticus* genome (GCA_001700915.1), hereafter called PasserD (PasserD.gff; Supplemental file 2), and compared it to the previous annotation, called Ensembl. The reannotation process (Figure 1) yielded a final set of 15,614 protein coding genes and 38,592 transcripts, resulting in an average of 2.6 transcript isoforms per gene at the whole-genome scale. In the PasserD annotation, 37,179 transcripts were found to have 3’UTRs or 5’UTRs, representing approximately 96.3% of all the annotated transcripts. The mean number of exons per gene increased from 12.5 in the prior annotation to 15.3 in PasserD. The PasserD annotation also contains alternatively spliced or alternatively initiated transcripts. Of the total transcripts, 87.2% (33,660) were assigned GO terms and 94.8% (36,567) were assigned complementary functional annotation, whereas these attributes were absent in Ensembl (Table 1).

**Table 1.** Summary of Ensembl and PasserD annotations.

| Type | Ensembl | PasserD |
| --- | --- | --- |
| **Protein-coding genes** |  |  |
| Number of genes | 14,188 | 15,614 |
| Number of transcripts | 22,926 | 38,592 |
| Number of exons | 291,615 | 641,666 |
| Transcripts with a 5' UTR | 10,379 | 31,917 |
| Transcripts with a 3' UTR | 11,147 | 36,234 |
| Transcripts with 5' and 3' UTR | NA | 28,128 |
| Mean gene length (bp) | 26,244 | 36,368 |
| Mean transcript length (bp) | 33,393 | 53,729 |
| Mean number of transcripts per gene | 1.6 | 2.6 |
| Mean CDS length (bp) | 1,987 | 2,467 |
| Mean number of exons per CDS | 12.5 | 15.3 |
| Mean exon length (bp) | 192 | 355 |
| Mean 5' UTR length (bp) | 270 | 784 |
| Mean 3' UTR length (bp) | 698 | 2,970 |
| Transcripts with associated GO terms | 0 | 33,660 |
| Transcripts with associated gene name | 0 | 36,525 |
| Transcripts with functional annotation | 0 | 36,567 |
| Complete BUSCOs | 89.20 % | 96.40 % |
| Missing BUSCOs | 7.80 % | 2.30 % |

### Evaluation and assessment of assembly completeness

We ran BUSCO in genome mode on the whole genome (GCA_001700915.1), obtaining 95.2% of completeness of the Passeriformes reference gene set, 6.8 % more than the benchmark obtained in Ensembl annotation (Figure 2).

**Figure 2.**
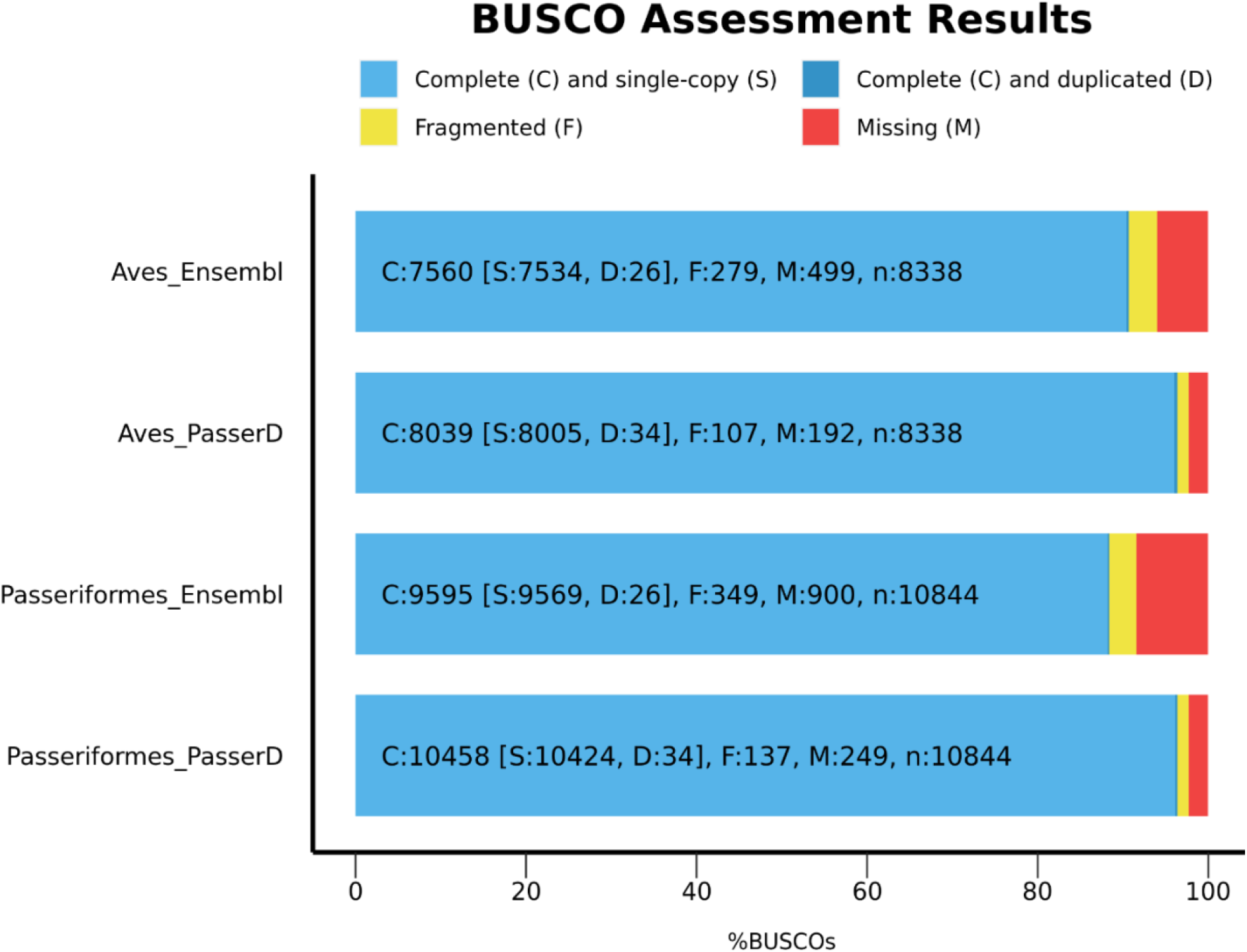
Comparison of gene set completeness between PasserD and Ensembl annotations of *Passer domesticus* genome. BUSCO scores were calculated by BUSCO 5.2.2 (*30*) in transcriptome mode using Passeriformes and Aves odb10 datasets.

PasserD reannotation and previous Ensembl annotation harbor 96.4% and 88.4% respectively of the 10,844 BUSCO Passeriformes lineage conserved genes being tested in transcriptome mode, or 96.4% and 90.7% respectively as well of the 8,338 BUSCO Avian conserved gene set (Figure 2).

### Prediction of gene functions

EggNOG assigned 33,469 transcripts to a specific GO term in PasserD reannotation (Supplemental file 3), a missed attribute in the previous annotation.

The improved accuracy of the new reannotation, over the previous, are illustrated in the following two examples. The first example deals with the PASSERD08661 gene, which encodes SI (sucrase-isomaltase enzyme), a protein expressed at the luminal side of the intestinal brush border membrane. This enzyme is essential to digest dietary carbohydrates including starch derivatives as maltose, isomaltose and malto-oligosaccharides, and sucrose (*33, 34*). SI in PasserD has extended evidence of additional exons and splicing sites with the integrations of 2 previously split genes in a new fully annotated gene with a longer transcript length (Figure 3a). The second example deals with the gene PASSERD06152, which was annotated as two separate genes in Ensembl whereas in PasserD it is a single gene. This gene encodes MUC2, a member of the mucin protein family, which are high molecular weight glycoproteins produced by many epithelial tissues. The protein encoded by this gene is secreted and forms an insoluble mucous barrier that protects the gut lumen (*35*) (Figure 3b). In both examples a gain of UTR regions can be recognized.

**Figure 3.**
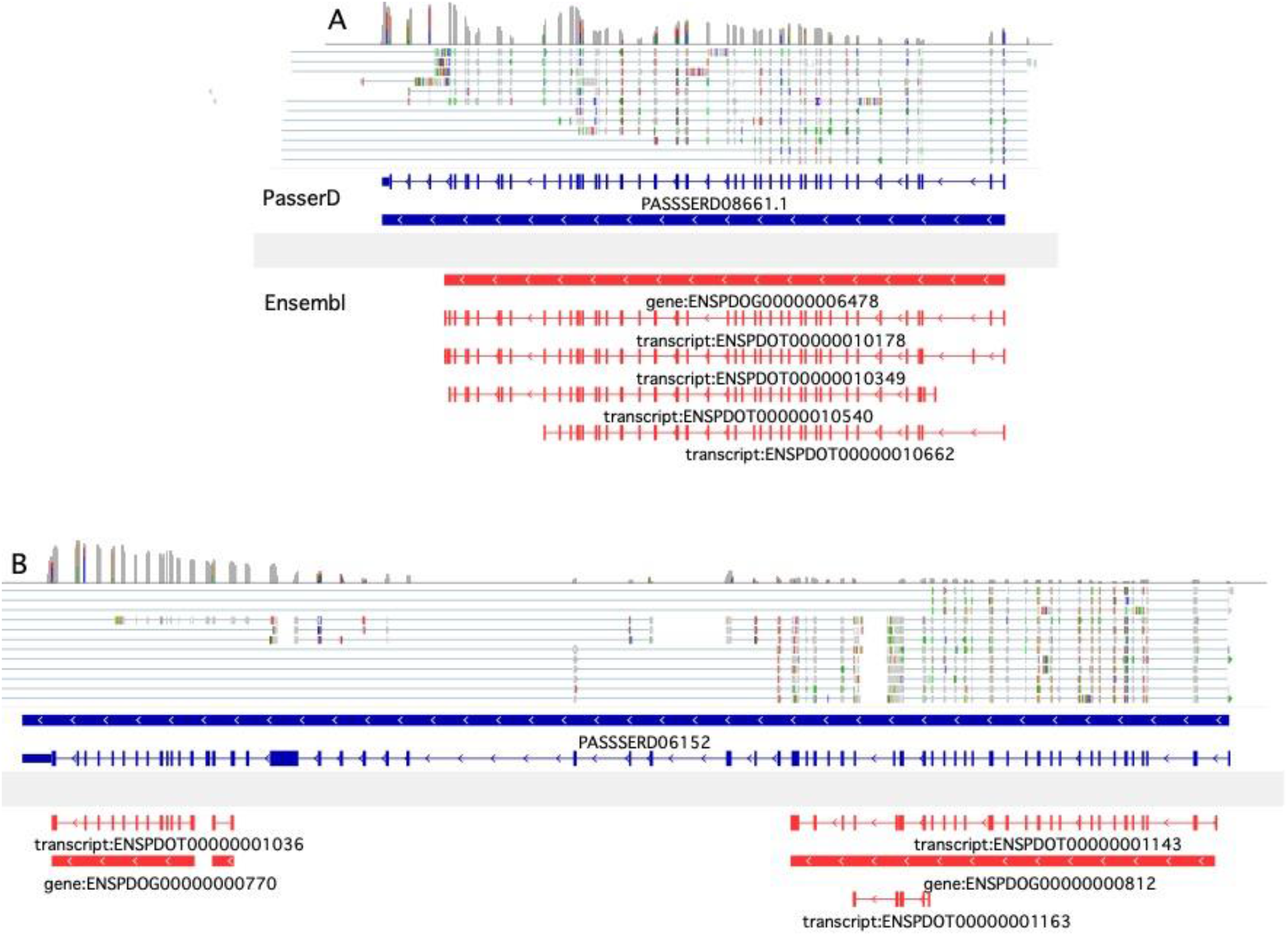
Examples of known genes with improved annotation. IGV views of the genes models of A) SI, B) MUC2 according to PasserD (blue) and Ensembl (red) annotations with RNA-seq mapped reads. The blue and red lines with gene IDs below represent the whole gene. The blue and red lines with mRNA IDs below show the exon and intron gene structure in detail. The thinner bars at the transcript ends represent UTRs, the thick bars represent exons forming the CDS (Coding Sequence), and the thin lines represent the introns. The arrowheads within the lines indicate transcriptional orientation.

Furthermore, those complete or fragmented genes missed in the previous annotation Ensembl were identified and their GO terms of the Biological Process domain analyzed using the overrepresentation test in Panther 17.0 (*36*). The results highlight that the pathways overrepresented by these genes are related to cellular processes involved in the assembly, arrangement of the constituent parts, or disassembly of a cilium, a filiform extrusion of the cell surface present in eukaryotic cells. This result is consistent with the fact that the included RNA-Seq data are of small intestinal origin, and it is expected to improve the gene annotation of those genes expressed exclusively and/or predominantly in this tissue. As in mammals, the intestinal epithelium in birds is a single layer of cells, mainly comprised of columnar absorptive cells (enterocytes) (*37*). In such cells, the apical surface area is greatly increased by microvilli, tightly packed finger-like projections of the membrane into the lumen. These microvilli may be considered as the primary cellular interface between the lumen and the inside of the organism in vertebrates, housing numerous proteins related to hydrolytic and absorptive processes. The exceptionally uniform size, orientation, and density of microvilli are generated by a complex network of proteins and signaling molecules, such as Villin and Plastin 1, which bundle actin filaments, in coordination with Ezrin and Myosin 1a (Myo1a) that crosslink the core actin bundles to their surrounding membrane (*38*). The genes for all these proteins are found in the new annotation PasserD (Table 2) and the improved annotation of these genes is depicted in Figure S1 (Supplemental file 1).

**Table 2.** Gene enrichment analysis of Gene Ontology terms of the Biological Process domain of the new genes annotated in PasserD using the overrepresentation test (Fisher test statistics) in PANTHER (*36*). From left to right columns contain: the name of the annotation data category, Ref.: number of genes in the reference list used (*Homo sapiens*), New: number of genes of the new annotation PasserD used, EXP: number of expected genes based on the reference list, f.e.: fold enrichment factor of the genes in the uploaded list over the expected, o/u: indicates if the category is over (+) or under (-) represented, *P*: raw value *P*-value determined by Fisher’s exact test (*P* < 0.05 is considered statistically significant), FDR: False discover rate (FDR) calculated by the Benjamini-Hochberg procedure (a critical value of 0.05 was used to filter results.

| GO biological process complete | Ref. | New | exp | f.e. | o/u | *P* | FDR |
| --- | --- | --- | --- | --- | --- | --- | --- |
| cilium flagellum assembly (GO:0120316) | 36 | 9 | 1.56 | 5.76 | + | 8.26E-05 | 3.01E-02 |
| cilium movement (GO:0003341) | 170 | 21 | 7.38 | 2.84 | + | 4.95E-05 | 2.16E-02 |
| microtubule-based process (GO:0007017) | 809 | 62 | 35.13 | 1.76 | + | 3.99E-05 | 2.02E-02 |
| motile cilium assembly (GO:0044458) | 62 | 12 | 2.69 | 4.46 | + | 5.10E-05 | 2.10E-02 |
| cilium assembly (GO:0060271) | 322 | 35 | 13.98 | 2.5 | + | 2.83E-06 | 4.44E-03 |
| organelle assembly (GO:0070925) | 803 | 61 | 34.87 | 1.75 | + | 5.56E-05 | 2.18E-02 |
| organelle organization (GO:0006996) | 3026 | 176 | 131.39 | 1.34 | + | 8.04E-05 | 3.00E-02 |
| plasma membrane bounded cell projection assembly (GO:0120031) | 418 | 40 | 18.15 | 2.2 | + | 1.35E-05 | 9.60E-03 |
| cell projection assembly (GO:0030031) | 430 | 41 | 18.67 | 2.2 | + | 1.03E-05 | 8.06E-03 |
| cilium organization (GO:0044782) | 355 | 38 | 15.41 | 2.47 | + | 2.25E-06 | 5.03E-03 |

As for computing resources, annotation pipelines demand a server grade computer, with a large amount of RAM and tens of CPU cores. An additional benefit obtained using the proposed pipeline is that all the required steps to prepare the data (e.g., Funannotate, Trimgalore, Hisat2 or Stringtie) can be run in a personal workstation (8-16 cores and 16-32GB of RAM) and then use a computational system with more resources to complete the GeMoMa step. De novo annotation of the house sparrow genome was completed in 4 h using 20 CPUs and the peak RAM usage was lower than 100GB (high end workstation or entry level server) and a load of 6 BAM files of 5 to 6 GB each. Even faster computational times can be achieved using additional CPUs in each step and run times are both a function of the evidence data set presented for alignment as well as the gene density of a genome, but the observed throughput of greater than 200 Mb h^−1^ demonstrates that even the largest of eukaryotic genomes could be annotated in a reasonable period of time.

## Conclusions

The genome annotation of protein coding genes is a necessary step for all downstream analyses and the choice of used resources and methods significantly impacts annotation quality and completeness.

Here, we used fast and low demanding computational resources, and an optimized annotation pipeline to improve the *P. domesticus* genome annotation by taking advantage of Illumina short reads obtained from intestinal tissue.

It not only updated the gene models, but also increased alternatively spliced isoforms and achieved a total of 80.9% and 93.11% of transcripts containing 5’ and 3’ UTRs, respectively. Given the material used, we extend the ongoing development of genomic resources for house sparrows focusing on functional genomics (*39*).

The house sparrow transcriptome data provided here can be used as a reference for comparative and differential expression analysis of gene expression, a significant and vibrant new opening venue for studying mechanistic bases of processes within and among species. Such comparisons provide a valuable resource for the advance of many fields including ecological and evolutionary physiology, conservation biology, and ecotoxicology. Therefore, its impact is likely to offer important insights to animal and human biomedical scientists.

## Supplemental Materials

***Supplemental file 1***. Figure S1. Examples of genes codifying structural microvilli proteins with improved annotation. Table S1. House sparrow intestinal RNASeq samples identification and ID accession numbers of this study

**Supplemental file 2**. PasserD.gff; gene annotation of the *P. domesticus* genome (GCA_001700915.1).

**Supplemental file 3**. New annotation PasserD. From left to right columns contain: Transcript name, Description of the transcript function, Preferred name of the transcript, GOs: GO term numbers, EC: Enzyme Commission number1, KO: KEGG Orthology Database code^2^, KEGG_Pathway code^2^, CAZy: Carbohydrate-Active EnZymes database code^3^, PFAM: database of protein domain families code^4^. References: ^1^ Bairoch (*40*), ^2^Kanehisa, Furumichi, Sato, Kawashima and Ishiguro-Watanabe (*41*), ^3^Cantarel, Coutinho, Rancurel, Bernard, Lombard and Henrissat (*42*), ^4^Sonnhammer, Eddy and Durbin (*43*).

## Acknowledgements

The UW Madison Center for High Throughput Computing (CHTC) is supported by UW-Madison, the Advanced Computing Initiative, the Wisconsin Alumni Research Foundation, the Wisconsin Institutes for Discovery, and the National Science Foundation, and is an active member of the OSG Consortium, which is supported by the National Science Foundation and the U.S. Department of Energy’s Office of Science.

