## Supplemental file 1 for "Rapid genome functional annotation pipeline anchored to the House sparrow (*Passer domesticus*, Linnaeus 1758) genome reannotation"

### Supplementary file 1

Figure S1. Examples of genes codifying estructural microvilli proteins with improved annotation. IGV views of the genes models of A) MYO1A and B) VIL1 according to PasserD (blue) and Ensembl (red) annotations with RNA-seq mapped reads. The blue and red lines with gene IDs below represent the whole gene. The blue and red lines with mRNA IDs below show the exon and intron gene structure in detail. The thinner bars at the transcript ends represent UTRs, the thick bars represent exons forming the CDS (CoDing Seguence), and the thin lines represent the introns. The arrowheads within the lines indicate transcriptional orientation.

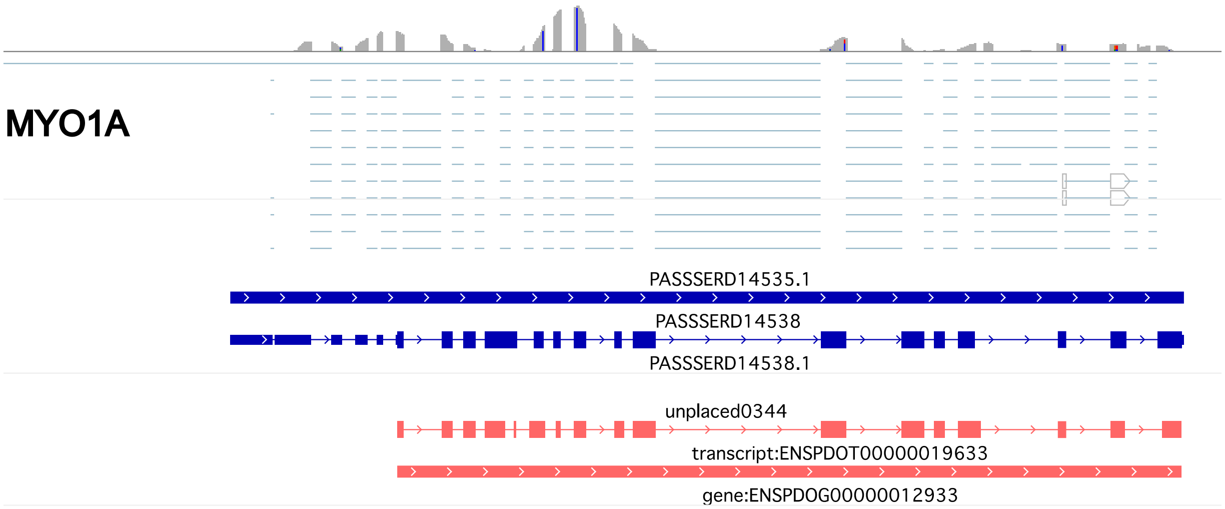


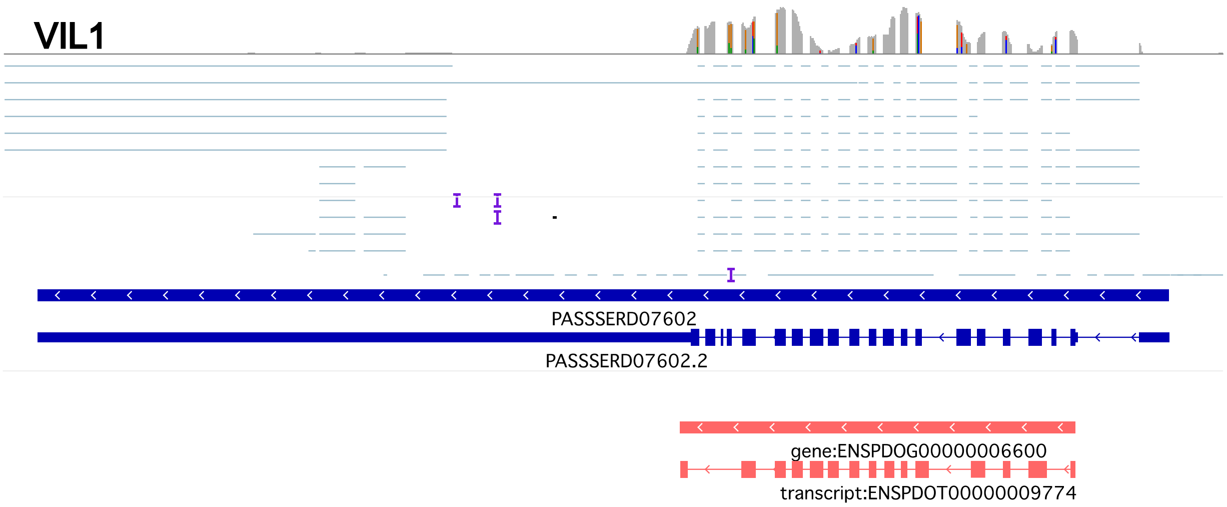

Table S1. House sparrow intestinal RNASeq samples identification and ID accession numbers of this study

| Bird | Sample | Access Id |
| --- | --- | --- |
| 527 | KV6 | SRS11440915 |
| 417 | KV32 | SRS11440916 |
| 433 | KV33 | SRS11440914 |
| 442 | KV34 | SRS11440913 |
| 471 | KV35 | SRS11440912 |
